# Multiple evolutionary events in host plant adaptation in Lepidoptera

**DOI:** 10.1101/2025.04.05.647383

**Authors:** Mei Luo, Bin Li, Lixin Ma, Zhongqiang Jia, Zhan Shi, Hangwei Liu, Zinan Wang, Bo Zhang, Sheng Yu, Jinfeng Qi, Yutao Xiao, Shaoqun Zhou, Henry Chung, Guirong Wang

**Author notes:** These authors contributed equally (Mei Luo, Bin Li, Lixin Ma).

## Abstract

The evolution of insect host adaptation is a key component of insect-plant coevolution, a complex process often shaped by multiple evolutionary events. In this study, we identified two UDP-glycosyltransferase (UGT) genes, *SfruUGT33T10* and *SfruUGT33F32*, in the fall armyworm *Spodoptera frugiperda*, which play critical roles in the tolerance of benzoxazinoids (BXs), secondary metabolites in maize. These two detoxification enzymes exhibited distinct glycosylation patterns for BXs and varying detoxification efficiencies, reflecting independent evolutionary trajectories. Phylogenetic analyses revealed that *SfruUGT33T10* originated independently within Noctuidae, while *SfruUGT33F32* resulted from tandem duplication within the UGT33F gene family and may have undergone neofunctionalization within the *Spodoptera* genus. Our findings provide evidence that the evolution of these two UGT paralogs contributed to the variation in the tolerance to maize BXs among different Lepidopteran species. This research underscores the significance of multiple independent evolutionary routes in host plant adaptation and offer new insights into the complex evolutionary processes underlying insect-plant interactions.

## Introduction

During the 300 million years of coevolution between herbivorous insects and their host plants, a continuous evolutionary arms race has been maintained (Ehrlich and Raven 1964, Guo et al. 2023). Plants have evolved diverse specialized metabolites to defend against herbivore attacks, which in turn drives herbivorous insects to evolve detoxification mechanisms as a counter-defense strategy to colonize their hosts (Heidel-Fischer and Vogel 2015, Erb and Kliebenstein 2020, Zhou and Jander 2021, Luo et al. 2023). The evolution of insect host adaptation is a key component of insect-plant coevolution, and gaining insight into this process is essential for uncovering the evolutionary forces that drive insect biodiversity and niche expansion (Després et al. 2007, Heidel-Fischer and Vogel 2015, Simon et al. 2015). Insect host adaptation is a complex and ongoing evolutionary process, typically shaped by multistep, multi-pathway adaptations (Futuyma and Agrawal 2009, Nylin and Janz 2009, Edger et al. 2015). However, current research primarily focuses on the functional analysis of single genes involved in adaptation, often neglecting broader evolutionary mechanisms.

Maize (*Zea mays*) is a major cultivated crop and a crucial food source worldwide. However, maize production is significantly challenged by pest infestations (Blanco et al. 2014). Among the various specialized metabolites produced by maize, benzoxazinoids (BXs) are particularly important due to their defensive roles against herbivores. BXs consist of more than 20 compounds that share a 2-hydroxy-2H-1,4-benzoxazin-3(4H)-one skeleton (Wouters et al. 2016, Zhou et al. 2018). The BXs biosynthetic pathway has been elucidated in maize, with the initial step catalyzed by the BX1 enzyme, which converts indole-3-glycerol phosphate into indole. Four cytochrome P450-dependent monooxygenases (BX2-BX5) subsequently mediate the stepwise oxidation of indoles, ultimately producing the toxic intermediate 2,4-dihy-droxy-1,4-benzoxazin-3-one (DIBOA). Through further enzymatic modifications, this pathway yields a series of benzoxazinoid glucosides, including DIBOA-Glc, DIMBOA-Glc (2,4-dihydroxy-7-methoxy-1,4-benzoxazin-3-one glucoside), and HDMBOA-Glc (2-hydroxy-4,7-dimethoxy-1,4-benzoxazin-3-one glucoside) (Frey et al. 1997, Tzin et al. 2017, Zhou et al. 2018). These compounds are stored as stable glucosides within plant cell vacuoles. Upon insect or pathogen attack, these glucosides are hydrolyzed by β-glucosidases, resulting in the release of toxic aglucones (Wouters et al. 2016). Among the most well-known toxic BX aglucone is DIMBOA, which has been shown to inhibit growth and reduce survival rates of various insect species, including *Ostrinia furnacalis* (Yan et al. 1999), *Spodoptera exigua* (Rostás 2007) and the aphid *Rhopalosiphum padi* (Mukanganyama et al. 2003).

Extensive research over the past two decades has identified several insect gene families, including cytochrome P450s (P450s), glutathione-S-transferases (GSTs), and UDP-glycosyltransferases (UGTs), as key players in the detoxification of plant secondary metabolites (Heidel-Fischer and Vogel 2015, Luo et al. 2023). Notably, UGTs have been recognized as the primary enzymes responsible for the degradation of BX compounds, rather than P450s or GSTs (Kojima et al. 2010). The detoxification mechanism involves UGT-mediated glycosylation of toxic BXs in the insect gut, producing epimeric glucosides that are resistant to plant glucosidases, thereby preventing the conversion of these compounds into toxins (Wouters et al. 2014). Specifically, a UGT, *SfUGT33F28*, has been identified as a candidate gene involved in the re-glucosylation of DIMBOA in the gut of the fall armyworm *Spodoptera frugiperda* (Israni et al. 2020, Israni et al. 2022). The elucidation of BXs detoxification mechanisms in Lepidopteran insects, along with the identification of the diverse BX compounds in maize, provides an excellent model for studying the complex processes of insect host adaptation and evolutionary dynamics.

In this study, we uncovered two independent evolutionary adaptations in Lepidoptera species that enable them to tolerate BX and feed on maize. Focusing on *S. frugiperda*, we identified two UGT genes, *SfruUGT33T10* (*Sfru33T10*) and *SfruUGT33F32* (*Sfru33F32*), involved in BXs detoxification. Our results showed that *Sfru33T10* originated and independently evolved within the Noctuidae family, conferring only limited effects on BXs detoxification. In contrast, *Sfru33F32* evolved specifically within the *Spodoptera* genus and confers higher tolerance to BXs, likely contributing to the successful colonization of maize by *Spodoptera* species. Our study highlights the importance of multiple independent evolutionary adaptations in Lepidoptera for overcoming plant chemical defenses, specifically against BXs in maize.

## Results

### Different effects of maize benzoxazinoids (BXs) on larval growth among five Lepidopteran species

To investigate the tolerance of Lepidopteran species on maize benzoxazinoids (BXs), we compared the weight growth of the newly hatched larvae of five Lepidopteran species (*S. frugiperda*, *S. litura*, *Mythimna separata*, *Helicoverpa armigera,* and *O. furnacalis*) when feeding on wild-type maize (W22) and on maize deficient in BX biosynthesis (bx2::Ds) (**Fig 1A**). The bx2::Ds line is a line with the BX2 gene knocked out, which leads to only trace amounts of BX produced, effectively rendering it a BX-deficient maize (Tzin et al. 2017) (**S1 Fig**). We hypothesized that species that cannot tolerate maize BXs could have better growth when feeding on BX-deficient maize. Our experiments showed that there was no significant difference in weight after seven days between larvae fed on W22 and bx2::Ds maize lines for *S. frugiperda* (*p* = 0.852) and *S. litura* (*p* = 0.390), while the larvae of *M. separata* (*p* = 0.003), *H. armigera* (*p* = 0.007), and *O. furnacalis* (*p* < 0. 001) had higher body weight when feeding on bx2::Ds maize (**Figs 1B-F**). These results suggest *S. frugiperda* and *S. litura* can tolerate the presence of BXs in maize, but growth of the other three Lepidopteran species is affected by BXs.

**Fig 1.**
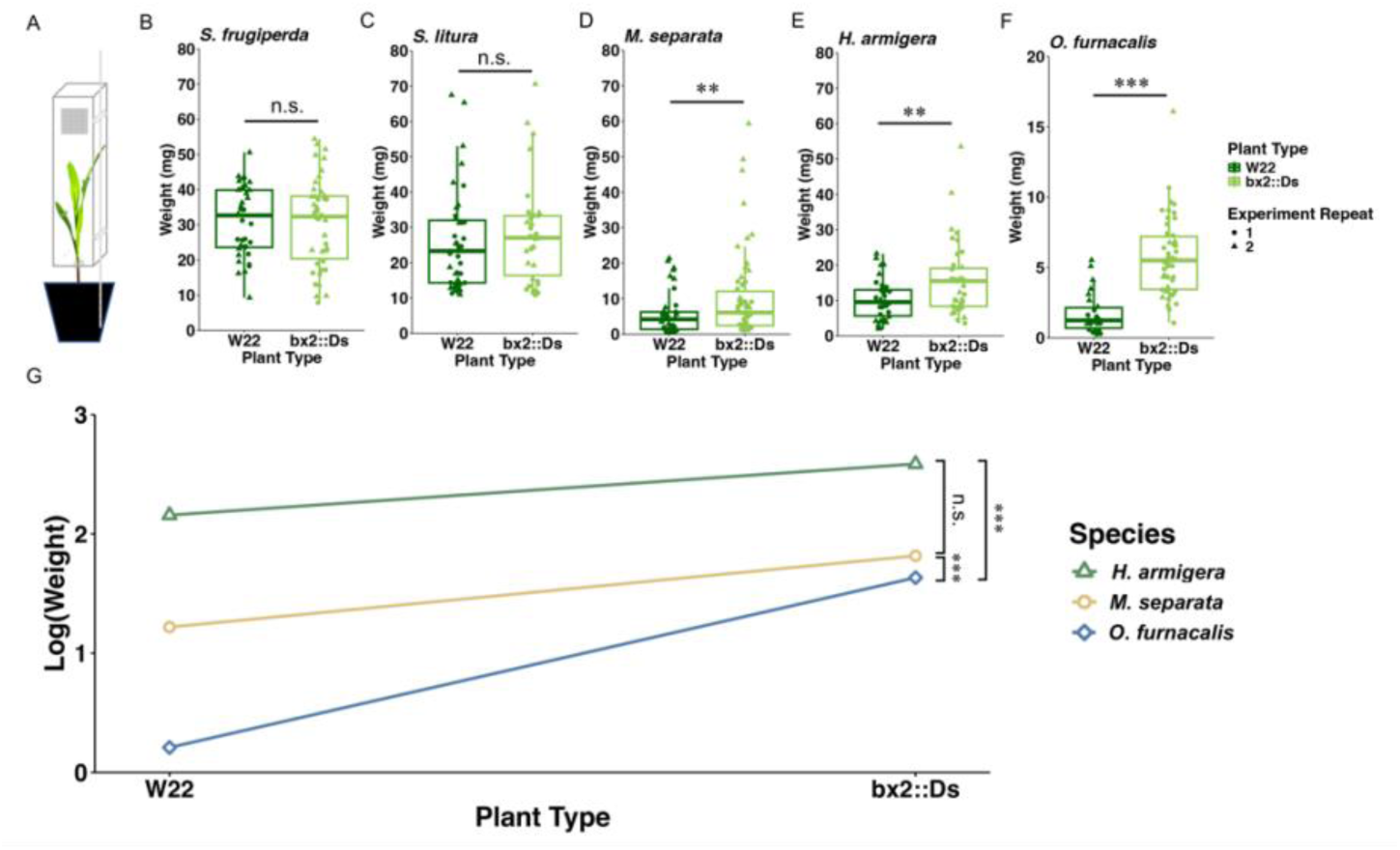
Differential tolerance of Lepidopteran species to maize benzoxazinoids (BXs). (A) Schematic diagram of feeding bioassay. The effects of feeding on wild-type W22 and bx2::Ds mutant maize on the larval weight of (B) *S. frugiperda*, (C) *S. litura*, (D) *M. separata*, (E) *H. armigera* and (F) *O. furnacalis*. The newly hatched larvae of different insects were inoculated onto two maize lines and weighed after 7 days of feeding. For each species, the data analysis was performed to compare insect weight across different plants. A linear mixed-effects model was constructed with plant type as a fixed effect and experiment repeat as a random effect to account for variation across experimental replicates. Boxplots were generated using ggplot2, with jittered scatter points overlaid to illustrate individual data distributions and experiment repeat-specific variation. The results were annotated with the corresponding *p*-value (α = 0.05). n.s. indicates nonsignificant differences, while **, and *** represent *p* < 0.01, and *p* < 0.001, respectively. (G) Effect of maize type (W22 vs. bx2::Ds) on larval weight gain of three Lepidopteran species. The body weights of these three Lepidopteran larvae were log-transformed and the differences in the larval body weight between treatments can be considered as the ratio of the changes. A generalized linear model was used to compare the larval weight gain across these Lepidopteran species, with insects species and plant types as fixed effects and the log-transformed body weight as the dependent variable. *P*-values were adjusted using the Benjamini-Hochberg correction (α = 0.05) in the custom. n.s. indicates nonsignificant differences, while *** represent *p* < 0.001, respectively.

We further asked whether there are differences in BX tolerance across three Lepidopteran species that exhibit incomplete tolerance to BXs. We fitted the body weight differences of the experiments in a generalized linear model and used customized contrasts to compare the body weight differences of these three species (**Fig 1G**). The contrasts showed that the body weight difference of *O. furnacalis* feeding on two maize lines was significantly higher than that of *M. separata* (*p* < 0.001) and *H. armigera* (*p* < 0.001) of which the larvae have increased body weights when feeding on bx2::Ds maize (**Fig 1G**). This suggests that *O. furnacalis* has lower tolerance than *M. separata* and *H. armigera*. Taking both analyses together, we suggest that there are differential levels of tolerance to BXs among these five Lepidopteran species.

### Two UGT genes contribute to the metabolism of BXs in *S. frugiperda*

As there were no differences in larval growth in *S. frugiperda* that are fed on W22 and bx2::Ds maize, we hypothesized that *S. frugiperda* exhibits a higher tolerance or enhanced ability to metabolize BXs compared to other Lepidopteran species. To determine the molecular mechanisms underlying BXs tolerance or metabolism in *S. frugiperda*, we conducted transcriptome sequencing (RNA-Seq) of midgut tissues from larvae fed on artificial diets (AF) without BX or wild-type maize (W22) for 24 hours. Principal component analysis (PCA) showed distinct expression profiles between the midgut expressed transcripts of AF- and W22-fed larvae (**S2A Fig**). As UGT genes have previously been implicated in the inactivation of BXs (Kojima et al. 2010, Wouters et al. 2014), we focused on gene expression differences in this gene family between AF- and W22-fed larvae in our transcriptomic analysis (**S2B Fig**). While many UGT genes had low or negligible expression in the midgut, a subset of UGT genes showed significantly higher expression in W22-fed larvae compared to AF-fed larvae (**Fig 2A and S1 Table**).

**Fig 2.**
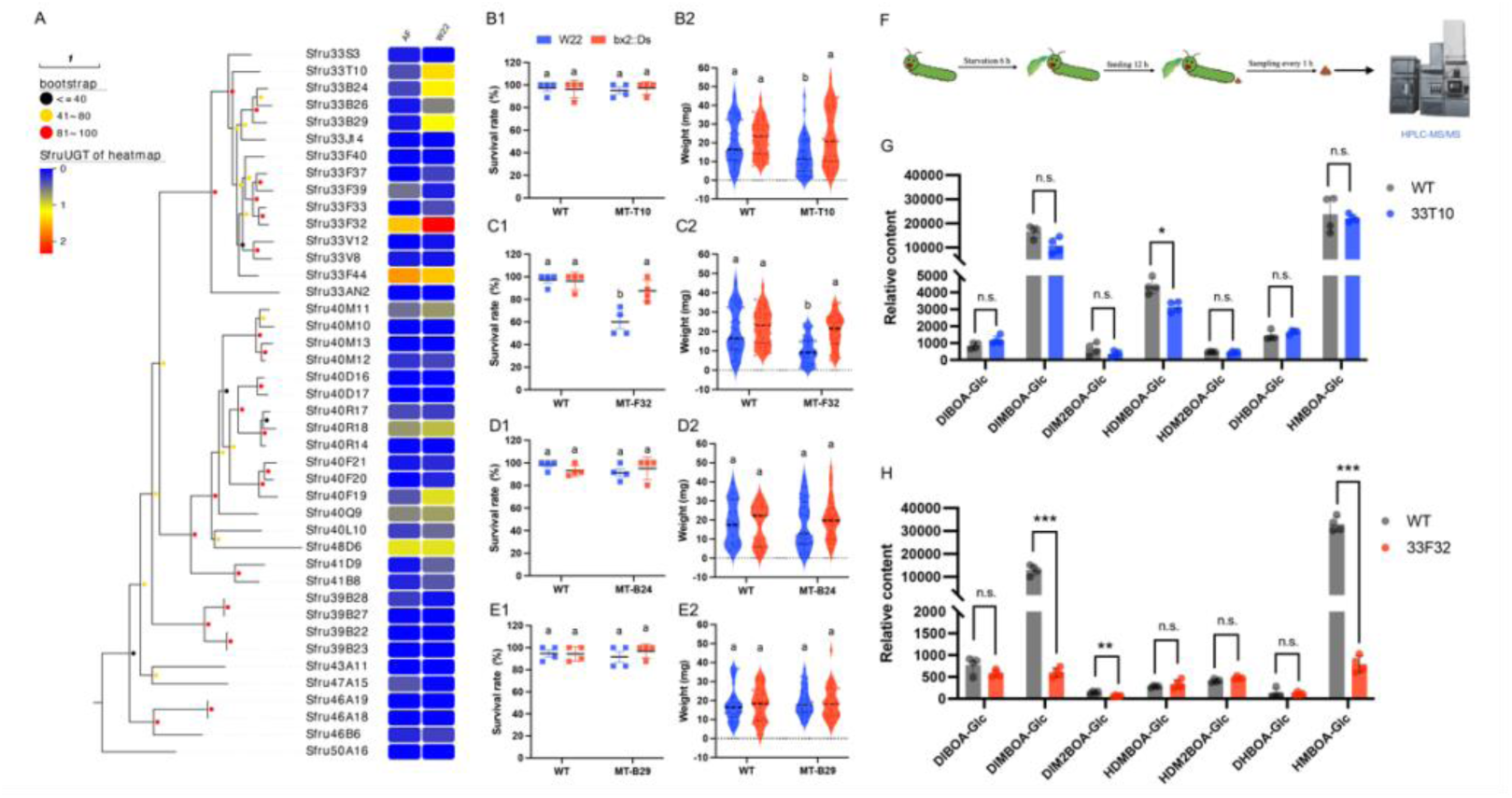
*Sfru33F32* and *Sfru33T10* are involved in the glycosylation detoxification of *S. frugiperda* to maize BXs. (A) Expression heatmap of SfruUGTs in *S. frugiperda*. The phylogenetic tree was constructed using maximum likelihood with 1000 bootstrap replicates. Samples were collected from the midguts of FAW that were fed on artificial food (AF) or wild-type maize (W22) for 24 hours, respectively. Expression levels of UGT genes were represented as log-transformed values, calculated as log_10_(FPKM + 1). The effects of feeding on wild-type W22 and bx2::Ds mutant maize on the survival rate (n = 4) and weight (n = 18 - 40) of (B) *Sfru33T10* mutants, (C) *Sfru33F32* mutants, (D) *Sfru33B24* mutants and (E) *Sfru33B29* mutants were investigated. Wild-type FAW was used as a control. Statistical analyses were performed using two-way ANOVA. Different letters represent significant differences at *α* < 0.05. (F) Schematic diagram of FAW larval feces sampling. Relative contents of Bxs in the feces of (G) *Sfru33T10* mutants and and (H) *Sfru33F32* mutants. The values represent the means ± SDs (n = 4). The relative content of each chemical was compared with that of the control to determine any significant differences using the Student’s *t*-test followed by Benjamini‒ Hochberg correction at *α* = 0.05. “n.s.” indicates nonsignificant differences; *, **, and *** indicate *p* < 0.05, *p* < 0.01, and *p* < 0.001, respectively.

We focused on four candidate UGT genes, *Sfru33T10*, *Sfru33F32*, *SfruUGT33B24* (*Sfru33B24*), and *SfruUGT33B29* (*Sfru33B24*), from the UGT33 subfamily, which displayed moderate to high expression (FPKM > 10, log_2_FoldChange > 2 and *p* < 0.05) (**S3 Table**) in the midgut. The differential expression of these genes was confirmed by qPCR (**S2C Fig**). To investigate the involvement of these candidate genes in BX metabolism, we generated homozygous knockout mutants of each of these genes via CRISPR/Cas9 (**S3 Fig**). We then compared the survival rate and larval weight of each of the mutant lines when they were fed on W22 and bx2::Ds maize. Among these four knockout lines, the knockout of *Sfru33F32* had the greatest effect on the ability of *S. frugiperda* to survive in W22 maize. *Sfru33F32* homozygous mutants exhibited a significantly lower survival rate and significantly lower larval weight when fed on maize W22 compared to bx2::Ds mutant maize, suggesting that *Sfru33F32* was involved in BX tolerance (**Fig 2C and S4B Fig**). The *Sfru33T10* homozygous mutant line, exhibited significantly lower larval weight when fed on maize W22 compared to bx2::Ds mutant maize, but the survival rate on the two maize types was not significant (**Fig 2B and S4A Fig**). This suggests that *Sfru33T10* may also contribute to BX tolerance but to a lesser extent than *Sfru33F32*. The survival rate and larval weight of *Sfru33B24* and *Sfru33B29* homozygous mutants did not show any significant differences in survival rate or larval weight when fed on W22 compared to bx2::Ds mutant maize, suggesting that these two UGT genes may not contribute to BX tolerance (**Figs 2D-E and S4C-D Fig**).

As UGTs are known to glycosylate toxic BXs into less harmful glycosides (Wouters et al. 2014), to determine whether *Sfru33T10* and *Sfru33F32* are involved in the metabolism of BXs, we quantified the relative amounts of seven BX glycosides in the fecal matter of both wild-type (WT) and mutant *S. frugiperda* larvae fed on maize W22 (**Fig 2F**). Compared with WT larvae, *Sfru33F32* mutants showed significantly reduced levels of three glycosylated compounds: DIMBOA-Glc (21.6-fold, *p* < 0.001), DIM_2_BOA-Glc (2.3-fold, *p* = 0.002) and HMBOA-Glc (41.7-fold, *p* < 0.001) (**Fig 2H**). Among all glycosylated compounds analyzed, only HDMBOA-Glc showed significantly lower levels in *Sfru33T10* mutants versus WT (*p* = 0.035) (**Fig 2G**). While both *Sfru33F32* and *Sfru33T10* contribute to the glycosylation and detoxification of BXs in *S. frugiperda*, they exhibited distinct glycosylation patterns for BXs and varying detoxification efficiencies.

### Evolution of *Sfru33F32* and *Sfru33T10* orthologs across Lepidoptera

To determine the evolutionary origins of *Sfru33F32* and *Sfru33T10* orthologs and variation in BXs tolerance among various Lepidopteran species, we conducted a comparative analysis of the UGT genes across key Lepidopteran species: five Noctuidae moths (*S. frugiperda*, *S. litura*, *S. exigua*, *M. separata*, and *H. armigera*), one Bombycidae species (*Bombyx mori*), two Pyralidae species (*Chilo suppressalis* and *O. furnacalis*), and one Plutellidae species (*Plutella xylostella*). All identified UGT genes were named according to the guidelines provided by the UGT Nomenclature Committee (**S2 Table**). Our results found that the UGT33 subfamily had the highest number of members across all species, followed by UGT40 (**S3 Table**).

Combined genome synteny (**Figs 3A1 and 3B1**) and phylogenetic analysis (**Fig 3C1 and S5 Fig**) showed that the homologs of *UGT33T10* exist in all five Noctuidae moths analyzed. In *S. frugiperda* and *H. armigera,* a single UGT33T gene exists at this locus, while the other two *Spodoptera* species, *S. litura* and *S. exigua*, contain three UGT33T genes at the syntenic locus. The UGT33T locus in *M. separata* contains four UGT33T genes. No corresponding syntenic loci for UGT33T genes are present in the genomes of *B. mori*, *C. suppressalis*, *O. furnacalis*, and *P. xylostella*, suggesting that UGT33T genes likely emerged after the divergence of Noctuidae from other Lepidoptera lineages. In contrast, there are significant tandem duplications at the UGT33F locus (**Fig 3B2**). There is a variable number of UGT genes at this locus found across the nine Lepidopteran species, including UGT33F homologs and other UGT genes. UGT33F subfamily genes are only found in the three *Spodoptera* species, *M. separata* and *H. armigera*. No UGT33F subfamily genes are found in this locus in *B. mori*, *C. suppressalis*, *O. furnacalis*, and *P. xylostella*. Despite the presence of other UGT genes at the UGT33F locus in these species, phylogenetic analysis reveals that these genes originated significantly earlier than the UGT33F subfamily (**S5 Fig**). To further confirm the orthologous relationships within the UGT33F subfamily, we combined phylogenetic results (**Fig 3C2**) and orthology analysis of UGT33F locus genes (**S4 Table**). We showed that *Sfru33F32* appears to be specific to the *Spodoptera* genus, with single-copy genes found in *S. frugiperda* and *S. litura*, and two copies identified in *S. exigua* (33F_3_OG0000000) (**S4 Table**), suggesting the orthologous group containing *Sfru33F32* appears to have evolved specifically within the *Spodoptera* genus. We inferred that UGT33F gene originated within the Noctuidae and subsequently underwent tandem duplication.

**Fig 3.**
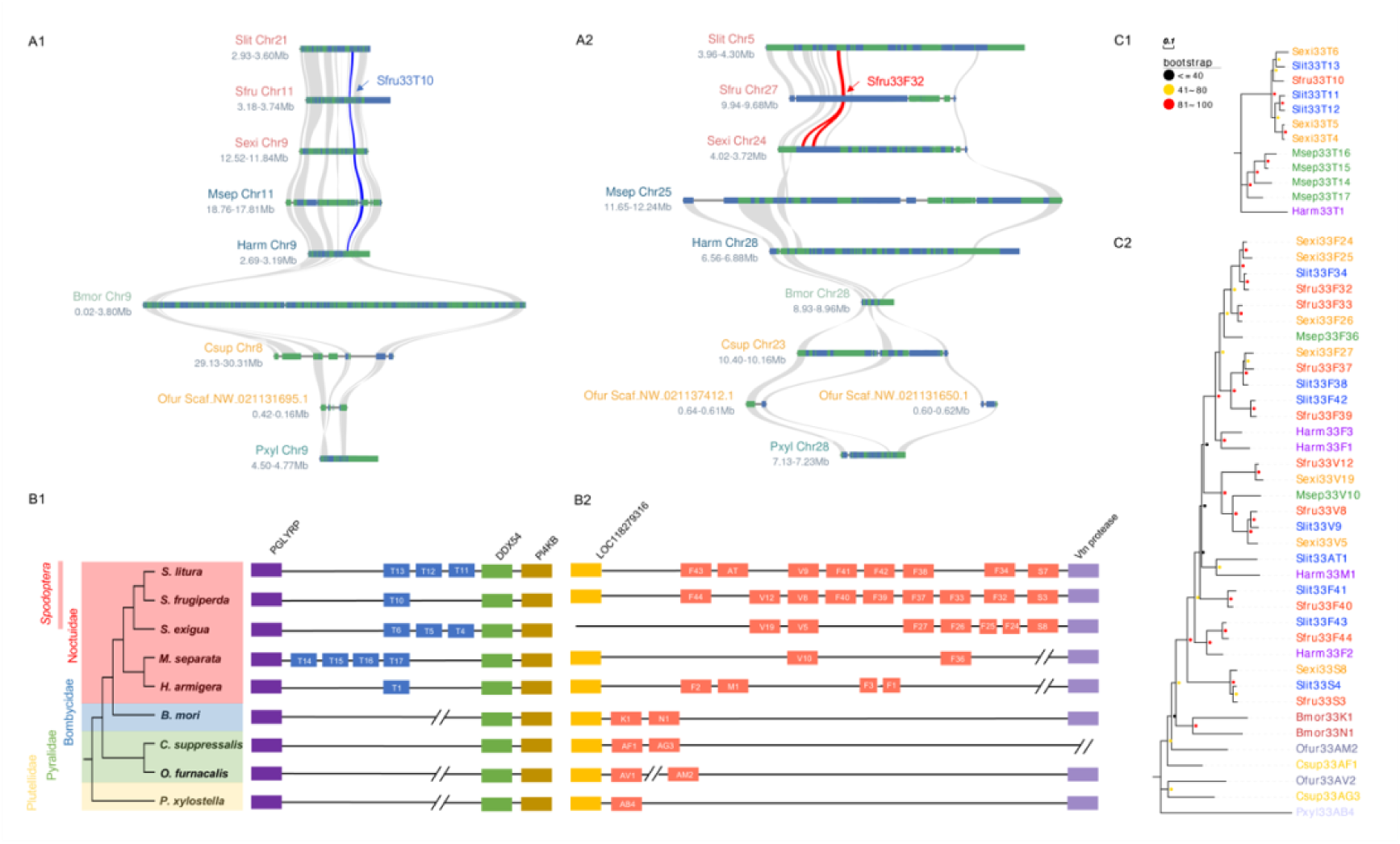
Evolutionary analysis of *Sfru33T10* and *Sfru33F32* orthologs across Lepidopterae. (A) Microsyntentic analysis of the (A1) *Sfru33T10* and (A1) *Sfru33F32* regions cross nine Lepidopteran species. The blue and red lines represent the syntenic regions of *Sfru33T10* and *Sfru33F32* and their orthologs in these species, respectively. The gray lines indicate gene collinearity across these nine species. (B) Gene duplication of the SfruUGT33 subfamily and the deep conservation across the nine lepidopteran species at the (B1) *Sfru33T10* and (B2) *Sfru33F32* loci. The phylogenetic tree was built with Orthofinder2 (version 2.5.4) using the longest transcripts of single-copy genes from nine Lepidopteran species. The blocks of red, blue, green and yellow cover five Noctuidae species, one Bombycidae species, two Pyralidae species and one Plutellidae specie in the tree, respectively. Orthologous genes or genes from the same UGT family have the same color block in the syntenic regions. (C) Phylogeny relationship of (C1) UGT33T and (C2) UGT33F block genes from 9 Lepidoptera species. The tree was constructed using maximum likelihood with 1000 bootstrap replicates. Bootstrap values are represented by circles at branch nodes, with the size and color of circles indicating support levels. The color of leaf labels represents different species.

### Evolution of BX tolerance across Lepidoptera

Our toxicology experiments showed differential levels of BXs tolerance across Lepidopteran species and knockout experiments in *S. frugiperda* showed the contribution of two genes, *Sfru33T10* and *Sfru33F32*, to BXs tolerance. While the knockout of *Sfru33F32* has a large effect on BXs tolerance in *S. frugiperda*, the knockout of *Sfru33T10* produces a smaller effect, likely due to the presence of *Sfru33F32*. Our evolutionary analysis showed that *Sfru33T10* orthologs are present in *M. separata* and *H. armigera*, but not in *O. furnacalis.* We hypothesize that *Sfru33T10* orthologs in *M. separata* and *H. armigera* may have a small contribution to BX tolerance in these species. To determine whether the function of *Sfru33T10* orthologs with regards to BXs tolerance is conserved across species, we performed molecular docking analysis on *Sfru33T10* and the UGT33T genes with the highest sequence similarity to *Sfru33T10* from four additional species to evaluate their potential to bind BX molecules (**Fig 3C1 and S5 Table**). The molecular docking models indicated that the binding pockets of these UGT33T genes share structural similarities, particularly in their interactions with the toxic maize compound HDMBOA, suggesting that these UGT33T genes may perform conserved detoxification functions across different Noctuidae species (**S6A Fig**). Additionally, multiple sequence alignment (**S6B Fig**) confirmed that several critical amino acid residues responsible for HDMBOA binding are conserved in *Sfru33T10* and its orthologs in other Noctuidae species, further supporting the hypothesis of conserved detoxification mechanisms. These findings suggest that the UGT33T subfamily may have retained a core functional role in detoxifying plant-derived compounds, such as BXs, across Noctuidae, with limited divergence in their binding mechanisms. This conservation likely reflects the evolutionary pressure to maintain effective detoxification capabilities in response to plant defenses.

The UGT33F locus contains multiple genes in addition to *UGT33F32* in *S. frugiperda*. To further investigate the role of other tandemly duplicated genes in the UGT33F region of *S. frugiperda*, we conducted functional validation of two additional genes, *Sfru33F37* and *Sfru33F44*. Using CRISPR/Cas9, we generated knockout mutants for each gene and assessed the impact of these mutations on maize BXs detoxification (**S7 Fig**). Survival rates and larval weights of the *Sfru33F37* and *Sfru33F44* mutants were measured when fed on both wild-type maize (W22) and the bx2::Ds mutant maize lacking BXs. The results showed that neither the survival rate nor the larval weight was significantly affected in the mutants compared to the wild-type controls (**Fig 4 A-B**), indicating that the knockout of *Sfru33F37* and *Sfru33F44* did not impair the ability of *S. frugiperda* to detoxify BXs. Furthermore, the fecal content analysis of BX metabolites revealed no significant differences in the relative levels of DIMBOA-Glc, HDMBOA-Glc, and other BX glycosides between the wild-type and the knockout mutants (**Fig 4C**). These results confirmed that, despite being part of the tandem duplication in the *UGT33F* region, *Sfru33F37* and *Sfru33F44* do not play a significant role in BX detoxification. Based on these evidences, we hypothesize that the lineage-specific evolution of *Sfru33f32* and its orthologs within the *Spodoptera* genus, along with their involvement in detoxifying BXs, may represent the neofunctionalization of UGT33 subfamily genes following gene duplication.

**Fig. 4.**
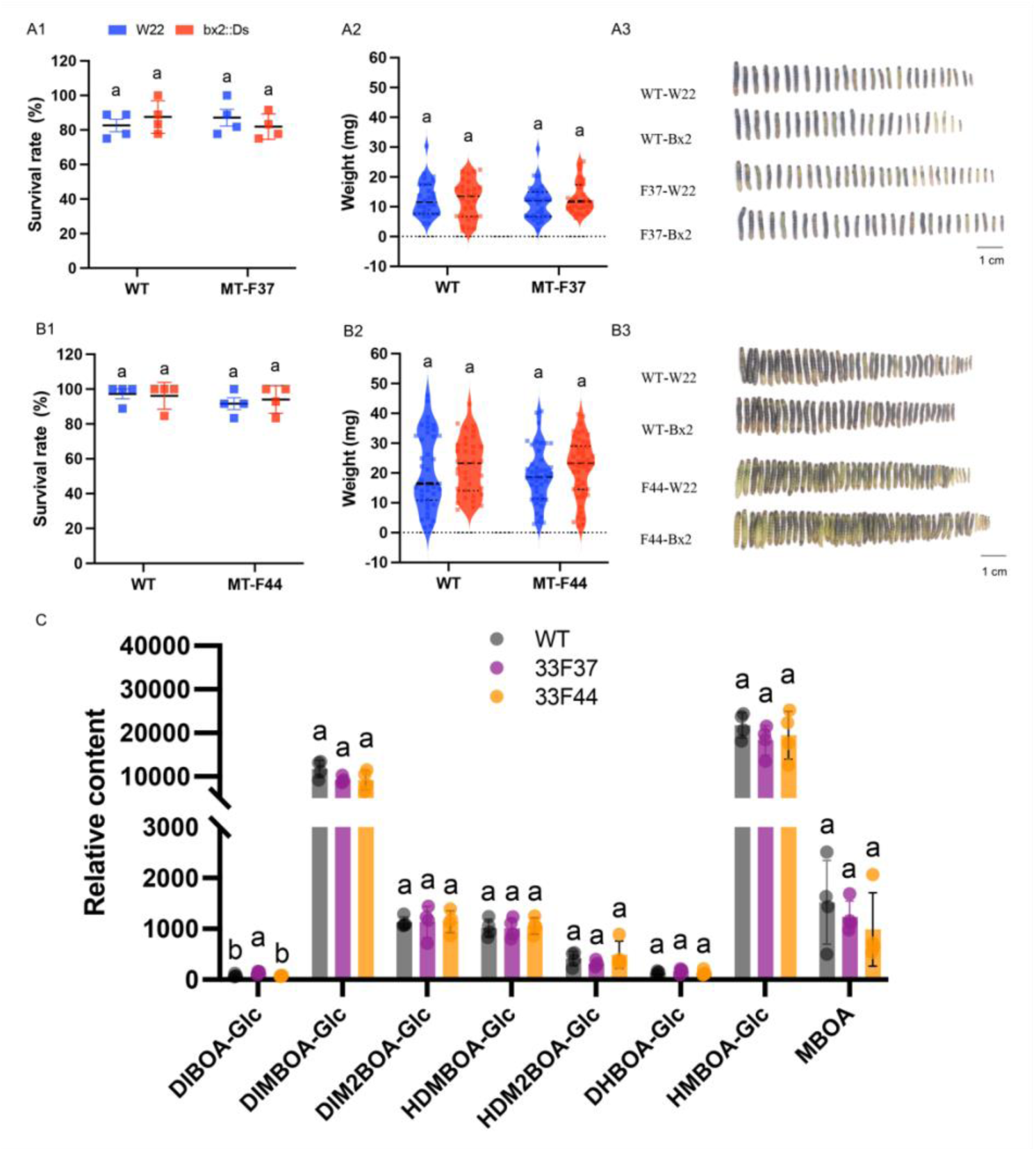
Sfru33F44 and Sfru33F37 do not participate in the detoxification of maize BXs in FAW. Effect of feeding on wild-type W22 and bx2::Ds mutant maize on the survival rate (n = 4), weight (n = 28-48) and larval picture of (A) *Sfru33F37* mutants, (B) *Sfru33F44* mutants. Statistical analyses were performed using two-way ANOVA. Different letters represent significant differences at alpha < 0.05. (C) The relative contents of Bxs in the feces of mutants of *Sfru33F37* and *Sfru33F44*. The values represent means ± SDs (n = 4). Different letters indicate significant differences within the same compounds as determined by one-way ANOVA (p < 0.05; post hoc tests).

## Discussion

In this study, we identified two key genes involved in the detoxification of BXs in *S. frugiperda, Sfru33T10* and *Sfru33F32*. *Sfru33F32* was found to be highly expressed in the midgut of *S. frugiperda* under both basal conditions and after maize induction, demonstrating strong tolerance toward BXs. Consistent with previous findings, *Sfru33F32* (previously annotated as *SfruUGT33F28*) has been established as a critical player in BX detoxification, with *in vitro* studies demonstrating its ability to glycosylate DIMBOA (Israni et al. 2020, Wang et al. 2024). However, the role of *Sfru33F32* in glycosylating other BX compounds remains unclear. This knowledge gap may stem from the inherent instability of certain BX aglycones, which rapidly degrade, complicating the use of stable substrates in enzymatic assays (Glauser et al. 2011). Our results show that loss of *Sfru33F32* suppresses glycosylation across multiple BXs, suggesting that *Sfru33F32* plays a broader role in BX detoxification than previously recognized. Notably, UGT genes are known for their substrate promiscuity, interacting with various plant secondary metabolites and xenobiotics (Pan et al. 2019, Snoeck et al. 2019), which adds complexity to understanding UGT-mediated detoxification pathways (Yan et al. 1999, Rostás 2007, Handrick et al. 2016, Snoeck et al. 2019).

In contrast, the role of Sfru33T10 in BX detoxification has received limited attention. Unlike Sfru33F32’s strong detoxification activity toward DIMBOA, Sfru33T10 exhibited a smaller effect on HDMBOA detoxification. The UGT33T subfamily has undergone species-specific tandem duplications in noctuid insects, with *S. frugiperda* and *H. armigera* possessing a single UGT33T gene, whereas other noctuids display varying levels of duplication. Despite its limited detoxification activity, Sfru33T10 shares conserved HDMBOA-binding sites with its orthologs in other species (**S6 Fig**), suggesting a potentially conserved role in BX detoxification. However, due to the rapid degradation of HDMBOA (Glauser et al. 2011), we were unable to purify stable HDMBOA for enzymatic assays, limiting our ability to directly assess UGT33T’s substrate-binding properties.

Gene expansion via large-scale duplication has been shown to contribute to host adaptation in herbivorous Lepidopterans (Fischer et al. 2008, Li et al. 2018, Heidel-Fischer et al. 2019). The UGT33 gene family exhibits notable expansion within Lepidoptera (**S2 Table**) (Ahn et al. 2012), with specific subfamilies, including UGT33F, displaying tandem duplications. In *S. frugiperda*, these duplicated UGT33F genes have undergone functional divergence; notably, *Sfru33F44* and *Sfru33F37* cannot detoxify BXs, while *Sfru33F32* has acquired this function. This finding aligns with recent studies demonstrating that, within the *S. frugiperda* UGT33F gene cluster, only *Sfru33F32* exhibits enzymatic activity toward DIMBOA, while the remaining four genes of UGT33F lack this capability (Wang et al. 2024). Moreover, our analysis reveals that the orthologous group containing *Sfru33F32* demonstrates lineage-specific evolution within the *Spodoptera* genus, suggesting that *Sfru33F32* and its orthologs likely have undergone neofunctionalization, thereby enhancing BXs detoxification in *Spodoptera* species compared to other Lepidopterans. These findings underscore the role of gene duplication in driving adaptive evolution in response to environmental pressures (Hahn et al. 2007).

We did not identify syntenic loci corresponding to *Sfru33T10* and *Sfru33F32* in the genome of *O. furnacalis*, a specialist maize pest, suggesting that the evolution of BXs tolerance in *O. furnacalis* occurred independently, potentially without reliance on the UGT33 gene family or through the involvement of other detoxification enzymes. From a host evolution perspective, *O. furnacalis* only began feeding on maize approximately the last 500 years, following the introduction of maize to Eurasia by Columbus (Calcagno et al. 2017). This relatively short evolutionary history of maize may explain its relatively low tolerance to BXs. Although *O. furnacalis* specializes in maize, it primarily feeds on stems and ears, which contain lower concentrations of BXs compared to roots and leaves (Davis et al. 2000, Oikawa et al. 2004).

In conclusion, we found that Lepidopteran insects that feed on maize exhibit varying degrees of adaptation to maize BXs. Specifically, *Spodoptera* species, such as *S. frugiperda* and *S. litura*, demonstrate greater BXs tolerance compared to other maize pests like *M. separata*, *H. armigera*, and *O. furnacalis*. In *S. frugiperda*, we identified two UGT genes, *Sfru33T10* and *Sfru33F3*2, that play key roles in BX detoxification. *Sfru33T10*, which evolved independently within noctuid insects, is involved in the detoxification of HDMBOA, providing a molecular basis for BX adaptation in Noctuidae. In contrast, *Sfru33F32* exhibited strong detoxification ability toward DIMBOA, HDMBOA, and DIM_2_BOA, contributing to the increased BX tolerance observed in *Spodoptera* species, which may result from the neofunctionalization of *Sfru33f32*. Our findings support the hypothesis that BX detoxification in Lepidoptera results from the multiple evolutionary trajectories of UGT genes. This study provides novel insights into the molecular mechanisms of host adaptation in major maize pests, including the economically significant *S. frugiperda*.

## Materials & methods

### Insect and plant materials

*S. frugiperda* (fall armyworms, FAW) were originally collected from maize fields in Yunnan Province, China, in 2019 and were reared in the laboratory for several generations under controlled experimental conditions. Eggs of *S. litura*, *M. separata*, *H. armigera,* and *O. furnacalis* were purchased from Keyun Biological Co., Ltd (Henan, China) and were also reared in the laboratory for more than two generations under controlled experimental conditions. The larvae of these species were reared on an artificial diet in an insect rearing chamber with a 14:10 light/dark photoperiod, a temperature of 25°C ± 2°C, and a relative humidity of 60% ± 5%. Adult moths were maintained under the same conditions and fed with a 10% sucrose solution after emergence.

Wild-type W22 and the bx2 mutant line Ds::bx2 were the *Z. mays* (maize) lines used in this study. Maize seedings were grown in 1-liter pots filled with a growing medium composed of a 5:1 mixture of potting soil and vermiculite. Plants were cultivated in a growth chamber with a 16:8 light/dark photoperiod and temperature set at 24 ± 2°C during the day and 20 ± 2°C at night. The plants were watered regularly and were used for experiments at the two-to three-week-old stage, with the second leaf fully developed and expanded from the whorl.

### Herbivore performance bioassay

To accurately measure the feeding performance of herbivores on maize, live plants were used instead of detached maize leaves in the experiments. To assess the adaptability of different herbivores to maize benzoxazinoids (BXs), five insect species that use maize as a host–*S. frugiperda*, *S. litura*, *M. separata*, *H. armigera*, and *O. furnacalis*–were tested to assess their adaptability to maize BXs. To prevent larval escape and ensure proper containment, a experimental setup was designed (**Fig. 1A**) Maize plants at the two– to three–week–old stage were enclosed in a perforated 8 × 8 × 30 cm^3^ (length × width × height) transparent PVC box. The base of each box was modified with a hole for the maize stem, which was wrapped in cotton and sealed with tape to prevent escape. For ventilation, rectangular openings (10 × 8 cm) were cut into two opposite sides of the box and covered with 200-mesh nylon screens. Newly hatched larvae were placed onto maize leaves within the box, and the top of the box was sealed with tape. Each box was supported vertically by inserting it onto a wooden stick that was anchored in the soil. Each box contained 3–4 maize plants, with 2–3 larvae per plant. After allowing the insects to feed freely on the maize for 7 days, the boxes were removed, and the number and weight of the herbivores were recorded. The insects were then frozen at −20°C for subsequent imaging. Each treatment included no fewer than 16 maize plants and 30 larvae.

### Gene knockout using CRISPR/Cas9

The construction of UGT33 mutants was based on the methodology outlined in previous research (Guo et al. 2020). To achieve a large-fragment deletion in the target UGT33 gene, two sgRNAs were designed to target distinct sites within the first exon, which is the longest exon of the four exons in UGT33. Initially, polymorphism detection was conducted on this exon to ensure a precise target design. We sequenced this region in ten female and ten male insects using the sequencing primers listed in **Table S7**. Genomic DNA from midlegs was extracted using the Multisource Genomic DNA Miniprep Kit (Axygen, New York, USA) according to the manufacturer’s recommendations. The PCR amplification was performed as follows: 98°C for 3 min; 35 cycles of 98°C for 10 s, 15 s at an annealing temperature appropriate for the designed primers, 72°C for 30 s; and 72°C for 10 min. The target design was performed using the online tool CRISPOR (http://crispor.org) (Concordet and Haeussler 2018). The sgRNAs were synthesized and assembled using the Precision gRNA Synthesis Kit (ThermoFisher Scientific, Catalog Number: A29377). sgRNA templates were synthesized via PCR. First, forward and reverse oligonucleotides for each sgRNA were synthesized using the Tracr Fragment + T7 Primer Mix to generate the DNA templates. The forward strand oligonucleotide consisted of a universal forward primer (5’-TAATACGACTCACTATAG-3’) and a gene-specific target oligonucleotide. In addition, the reverse strand oligonucleotide consisted of a universal reverse primer (5’-TTCTAGCTCTAAAAC-3’) and a gene-specific target reverse oligonucleotide. The primers used for template synthesis are listed in **S7 Table.** The synthesized template was then used for in vitro transcription to generate sgRNAs, which were subsequently purified using the components provided in Box 2 of the Precision gRNA Synthesis Kit.

The final sgRNA concentrations were adjusted to 500 ng/µL. The injection mixture (10 µL) was prepared by combining 4 µL of sgRNA1 (500 ng/µL), 4 µL of sgRNA2 (500 ng/µL), 1 µL of Cas9 protein (150 ng/µL), and 1 µL of phenol red (used to visualize the injected embryos). An ample number of FAW pupae were placed in insect rearing cages to allow for free eclosion and mating. Fresh embryos, less than 1 hour old, were collected for microinjection. The injection mixture was injected into preblastoderm embryos using a Drummond Nanoject III microinjector (Item No. 3-000-207) with pulled glass capillary needles (Sutter P-97). The injected embryos were allowed to develop to the G0 generation in an insect rearing chamber.

G0 adults were collected before mating, and midlegs were taken for genotyping. The gDNA extraction and PCR conditions were consistent with those used in the polymorphism detection section. PCR products were analyzed by agarose gel electrophoresis to identify individuals with two bands, indicating potential heterozygotes. These potential heterozygotes were further validated by Sanger sequencing to confirm the exact genotype and whether it resulted in a frameshift mutation. If no large-fragment deletions were found, individuals with single-target mutations causing frameshifts were retained. The identified heterozygotes were crossed with wild-type (WT) unmated adults to produce F1 individuals. F1 larvae were screened using the same methods to identify heterozygous mutants, which were then intercrossed to produce F2 individuals, expected to segregate with 25% homozygous mutants. These F2 homozygous individuals were subsequently bred to establish stable homozygous mutant lines for further experiments.

### Annotation of the UGT gene family

Although the UGT gene family have already been annotated in many insect species, to eliminate discrepancies arising from variations in genome quality and annotation methods, we reannotated the UGT genes in several species included in this study. Firstly, we constructed a local insect UGT database using UGT protein sequences from *Drosophila melanogaster* (Ahn and Marygold 2021), *B. mori* (Ahn et al. 2012), *H. armigera* (Ahn et al. 2012), *S. exigua* (Hu et al. 2019), *C. suppressalis* (Zhao et al. 2019), *P. xylostella* (Li et al. 2017), and *S. frugiperda* (Gouin et al. 2017). Using the high-quality genomes of *S. frugiperda*, *S. litura*, *S. exigua*, *M. separata*, *O. furnacalis* and *P. xylostella* as references, we performed a local BLASTP (Camacho et al. 2009) search between our database and these genomes. Furthermore, we utilized the previously obtained UGT dataset to construct a Hidden Markov Model (HMM) using the HMMER software (Finn et al. 2011). This model was subsequently employed to annotate the UGT sequences of the target insect species. The intersection of genes identified by both annotation methods was selected for further analysis.

The candidate UGT genes were manually annotated in several steps. First, in accordance with the UGT Nomenclature Committee’s guidelines on sequence length, sequences shorter than 350 amino acids and longer than 800 amino acids were removed. Subsequently, all genes were subjected to domain analysis using the InterPro website (https://www.ebi.ac.uk/interpro/) to verify domain structures, and signal peptides were predicted using SignalP 4.1 (https://services.healthtech.dtu.dk/services/SignalP-4.1/). Finally, all candidate genes were submitted to the UGT Nomenclature Committee for official naming. Genes were labeled with the species abbreviation followed by the gene name.

### Phylogenetic analysis

The phylogenetic analysis was conducted following our previous research (Luo et al. 2020, Zhang et al. 2023). For the phylogenetic analysis of the UGT gene family, amino acid sequences were aligned using MAFFT v7.505 (Katoh et al. 2019) and Maximum-likelihood searches were performed using RAxML v8.2.12 (Stamatakis 2014) with PROTGAMMALG model and 1000 bootstrap replicated. The number of UGT genes and their expression levels (heatmaps) were fitted and visualized alongside the phylogenetic tree using Evolview (Subramanian et al. 2019). A species phylogenetic tree was constructed using nine species. Orthofinder (version 2.5.4) (Emms and Kelly 2019) was employed with default settings to build the phylogenetic tree, utilizing the longest transcripts for each gene based on the genome database (**S6 Table**).

### Ortholog analysis

Orthologous relationships within the UGT33F locus were identified using OrthoFinder (version 2.5.4). Protein sequences corresponding to the UGT33F locus from five noctuid species were input for analysis. Sequence similarity searches were conducted with BLAST, followed by multiple sequence alignments using MAFFT. Maximum-likelihood phylogenetic trees were subsequently generated using RAxML-NG. Gene orthology inference was performed based on multiple sequence alignment (MSA), with an inflation value set to 4.

### Synteny analysis

The genomic information for the nine species is provided in Supplementary **S6 Table**, with phylogenetic tree construction following previously described methods. Whole-genome synteny across multiple species was detected and visualized using MCScanX (Python package, default parameters) (Wang et al. 2012). Syntenic blocks containing the target genes *Sfru33F32* and *Sfru33T10* were highlighted with specific color codes in the collinearity file. This visualization was combined with the phylogenetic tree of the UGT33 family to distinguish orthologs from paralogs, illustrating the correspondence of UGT gene positions across different species’ genomes.

### RNA-seq of larval midguts

To investigate the gene expression profiles of FAW under different feeding conditions, fourth-instar larvae of each species were fed fresh maize leaves (W22) for 24 hours. Larvae fed with artificial feed (AF) served as controls. After the feeding period, midguts were rapidly dissected and frozen in liquid nitrogen for further analysis. Each treatment consisted of three biological replicates, with five midguts pooled per replicate.

Total RNA was extracted from the midgut samples using the TRIzol reagent kit (Invitrogen, Waltham, MA, USA) according to the manufacturer’s instructions. RNA quality and concentration were assessed using a NanoDrop spectrophotometer, and RNA integrity was verified with an Agilent 2100 Bioanalyzer. RNA-Seq libraries were prepared using the NEBNext Ultra RNA Library Prep Kit (NEB, Ipswich, MA, USA) following the manufacturer’s protocol. Libraries were sequenced on an Illumina HiSeq 2500 platform to generate paired-end reads of 150 bp. After adapter sequences and low-quality reads were filtered out, the cleaned reads were mapped to the reference genomes of FAW and *S. litura* using the STAR 2.4.0 software (Dobin et al. 2013). Gene expression levels were quantified as fragments per kilobase of transcript per million mapped reads (FPKM). Gene model read counts were aggregated using the HTseq tool (Anders et al. 2015). Differentially expressed genes (DEGs) were identified using the DEseq2 package, with thresholds of FDR < 0.05 and FC ≥ 2 (Love et al. 2014). All processed reads are deposited in the NCBI online repository in fastq format and can be accessed under the project ID: GSE276946.

FPKM values for all genes were used to perform principal component analysis (PCA) in RStudio version 2021.09.1 (RStudio Team 2021). The FPKM values were normalized and log-transformed before analysis. The PCA was conducted using the prcom function in R, and the resulting principal components were plotted to illustrate the variance in gene expression between the two feeding treatments. The volcano plot was generated using TBtools (Chen et al. 2020), plotting the log2-fold change against the −log10 FDR to highlight significantly upregulated and downregulated genes between the AF and W22 conditions according to the DEG results. Expression levels of UGT genes were represented as log-transformed values, calculated as log10(FPKM + 1), to create the heatmap.

### Quantitative real-time PCR analysis

Total RNA was extracted using the Quick RNA Isolation Kit (Cat. No. 0416-50, Huayueyang, Beijing, China) according to the manufacturer’s instructions. cDNA was synthesized from 500 ng of RNA with the HiScript® III 1st Strand cDNA Synthesis Kit (Cat. No. R321-02, Vazyme, Nanjing, China). Quantitative real-time PCR (qRT‒PCR) was conducted on a QuantStudio Real-Time PCR System (Applied Biosystems, Waltham, MA) using SYBR qPCR Master Mix (Cat. No. Q711-02, Vazyme, Nanjing, China) in a final reaction volume of 20 μl. Gene expression levels were determined using the 2-(ΔΔCt) method. All sequences of primers used for qRT‒PCR are provided in Supplemental **S7 Table**.

### Benzoxazinoid extraction and quantification

FAW larvae were reared on an artificial diet until they reached the 4^th^ instar. Prior to being transferred to maize (W22), the larvae were starved for 6 hours. After being fed on maize for 12 hours, the maize leaves were replaced with fresh ones and insect frass was collected every hour. The collected frass was placed in preweighted 1.5 ml centrifuge tubes and freeze-dried at −80°C for 12 hours. After freeze-drying, the frass was weighed and stored in a −80°C freezer for future use. Approximately 20 ± 3 mg of each freeze-dried frass sample was used to extract BXs. The samples were suspended in MeOH/H2O (50:50, v/v; 0.5% formic acid) and were transferred to glass vials for analysis on an HPLC-MS/MS system (LCMS-8040, Shimadzu) with previously purified analytical standards according to Qi et al (Qi et al. 2016). The concentration of each BX was normalized to the frass fresh weight of the frass and the abundance of internal standards.

### Construction of protein models and molecular docking

The 3D structure models of *SlitUGT33T13*, *Sfru33T10*, *SexiUGT33T6*, *MsepUGT33T17* and *HarmUGT33T1* proteins were constructed using the AlphaFold 3 online server (Abramson et al. 2024). The 3D structure of HDMBOA (CAS: 149182-67-2) was obtained from the PubChem database (https://pubchem.ncbi.nlm.nih.gov/). The ligand docking pockets of the models were individually predicted using Fpocket (Le Guilloux et al. 2009, Schmidtke et al. 2010). The parameters of docking grid boxes of the were generated according to the ligand docking pockets using PyMOL 2.1 (Schrödinger, LLC, New York, NY, USA). For molecular docking, the hydrogens and charges of proteins and ligand chemicals were dealt with MGLTools 1.5.7 (Molecular Graphics Laboratory, La Jolla, CA, USA) to generate pdbqt files. The five protein models were docked with HDMBOA using AutoDock 4.2.6 (Morris et al. 2009), respectively. A total of 256 docking poses were generated in every docking process and were divided into several clusters by MGLTools 1.5.7. The protein-ligand interaction of the middle docking pose in the largest cluster was analyzed using the Protein-Ligand Interaction Profiler (PLIP) (Adasme et al. 2021). The complex structures and protein-ligand interactions were visualized and characterized using PyMOL 2.1 (Schrödinger, LLC). Multiple sequence alignments of *UGT33T* genes were performed using MAFFT, followed by the calculation of distance matrices using MEGA (version 10.2.6) to assess sequence divergence.

### Statistical analysis

To test whether the body weight differences between feeding treatments differed across five Lepidopteran species, the newly hatched larvae of different insects were inoculated onto two maize lines and weighed after 7 days of feeding. For each species, data analysis was performed to compare larvae weights across different plants. A linear mixed-effects model was constructed with plant type as a fixed effect and experiment repeat as a random effect to account for variation across experimental replicates. Boxplots were generated using ggplot2, with jittered scatter points overlaid to illustrate individual data distributions and experiment repeat-specific variation. The significance of the fixed effect was assessed using the *t*-test (*α* = 0.05) derived from the model summary. To test whether there are differences in BX tolerance across three Lepidopteran species that exhibit incomplete tolerance to BXs, we applied a generalized linear model, which includes the species, plant types, and their interaction as independent variables and the log-transformed body weight as the dependent variable. We then used customized contrasts to compare the body weight differences of the three species between species. *P*-values were adjusted using the Benjamini-Hochberg correction (*α* = 0.05) in the custom contrast. To compare BX content in the frass of *Sfru33F32* and *Sfru33T10* mutants following consumption of maize, the relative content of each chemical was compared with that of the control to determine any significant differences using the Student’s *t*-test followed by Benjamini‒Hochberg correction at *α* = 0.05. For the qPCR experiments, Student’s *t*-test was used to compare the expression levels between treatment and control groups. Single-factor pairwise comparisons were performed using Student’s *t*-test under the assumption of a normal distribution. For comparisons involving multiple samples with a single factor, one-way ANOVA was applied (compare BX content in the frass of *Sfru33F37* and *Sfru33F44* mutants following consumption of maize). For experiments assessing the effects of two factors, namely mutant compared to wild-type insects and maize variety, on survival rates and body weight, a two-way ANOVA was performed. When significant differences were found, *post hoc* analyses were conducted using Tukey’s test at *α* = 0.05. Statistical analyses and graphical representations were performed using Prism 9 software (GraphPad Software, San Diego, CA) and R (version 4.3.1).

## Acknowledgments

This work is supported by the National Natural Science Foundation of China (32202305 to M.L.), the China Postdoctoral Science Foundation (2022M723451 to M.L.), the Shenzhen Science and Technology Program (Grant No. KQTD20180411143628272 to G.W.), the Special Funds for Science Technology Innovation and Industrial Development of Shenzhen Dapeng New District (Grant No. PT202101-02 to G.W.), and the National Key Research and Development Program of China (2022YFD1700201 to G.W.). We thank Bin Gao, Peng Wang, Zhenxin Liu, Minghui Jin, and Zerui Deng for their assistance with insect rearing.

## Author Contributions

M.L., H.C. and G.W. conceived the overall research plan. M.L., S.Z. and G.W. designed the experiments. M.L., B.L., L.M., Z.S. and J.Q. performed the experiments. M.L. and Z.J. contributed to the data visualization. M.L., H.L., B.Z., Z.W. and S.Y. analyzed the data. G.W., S.Z. and Y.X. provided experimental platforms, germplasm resources, and other essential materials. M.L. and H.C. wrote the initial draft of the manuscript, and G.W. supervised the writing with input from all authors.

