## supplemental figure for "Multiple evolutionary events in host plant adaptation in Lepidoptera"

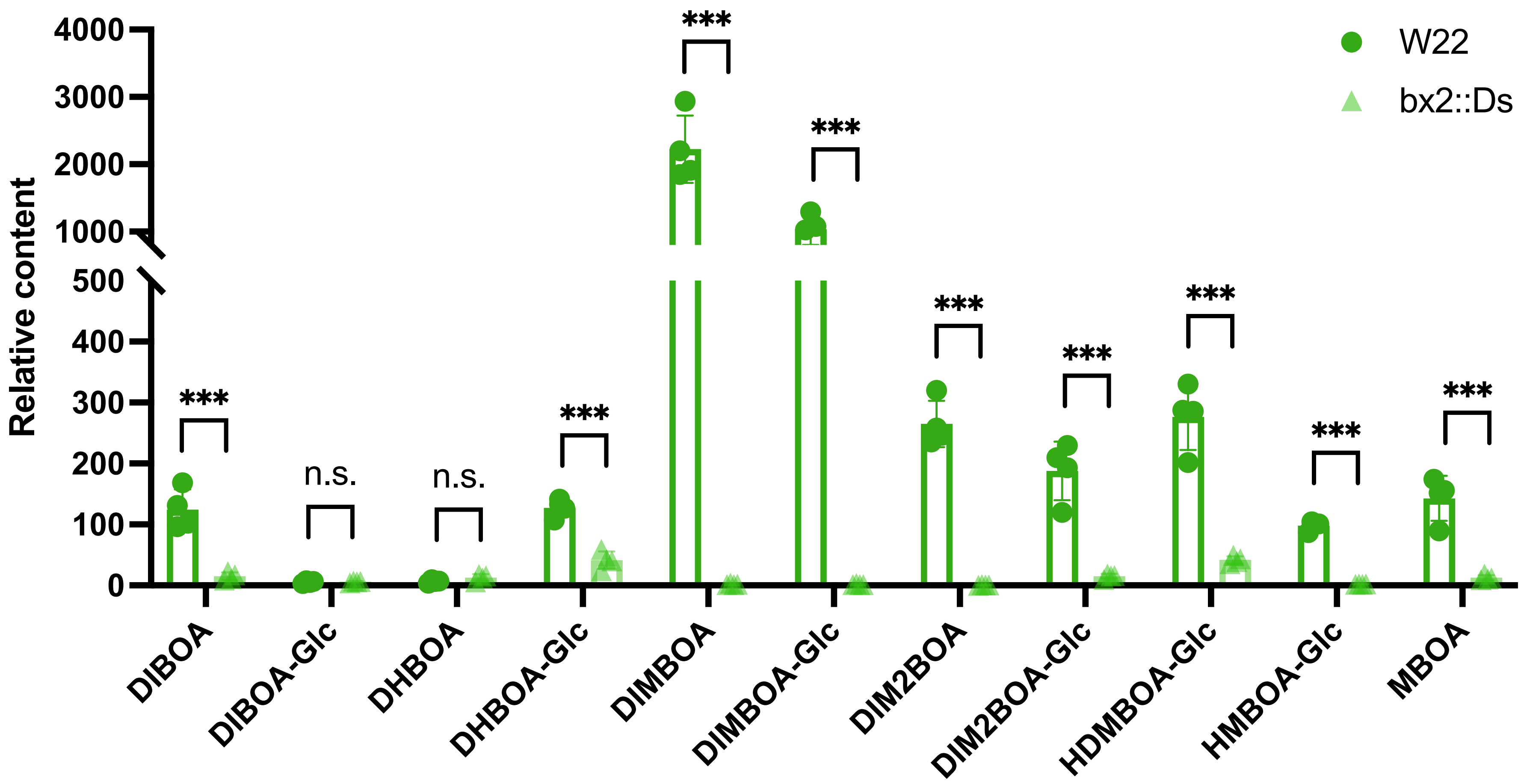


**Fig. S1. Relative contents of benzoxazinoids (BXs) in wild-type maize (W22) and bx2::Ds mutant plants.** W22 refers to wild-type maize, and bx2::Ds denotes maize line deficient in BXs. The contents of BXs were measured from the second leaves of two-week-old maize plants using high-performance liquid chromatography (HPLC). The data are presented as the means ± SDs (n=4). Differences in BX contents between maize lines were compared using Student's *t* test, and *p* values were adjusted for multiple comparisons using the Benjamini–Hochberg correction (α = 0.05). n.s. indicates nonsignificant differences, and *** indicates $p < 0.001$. Error bars represent standard error of the mean.

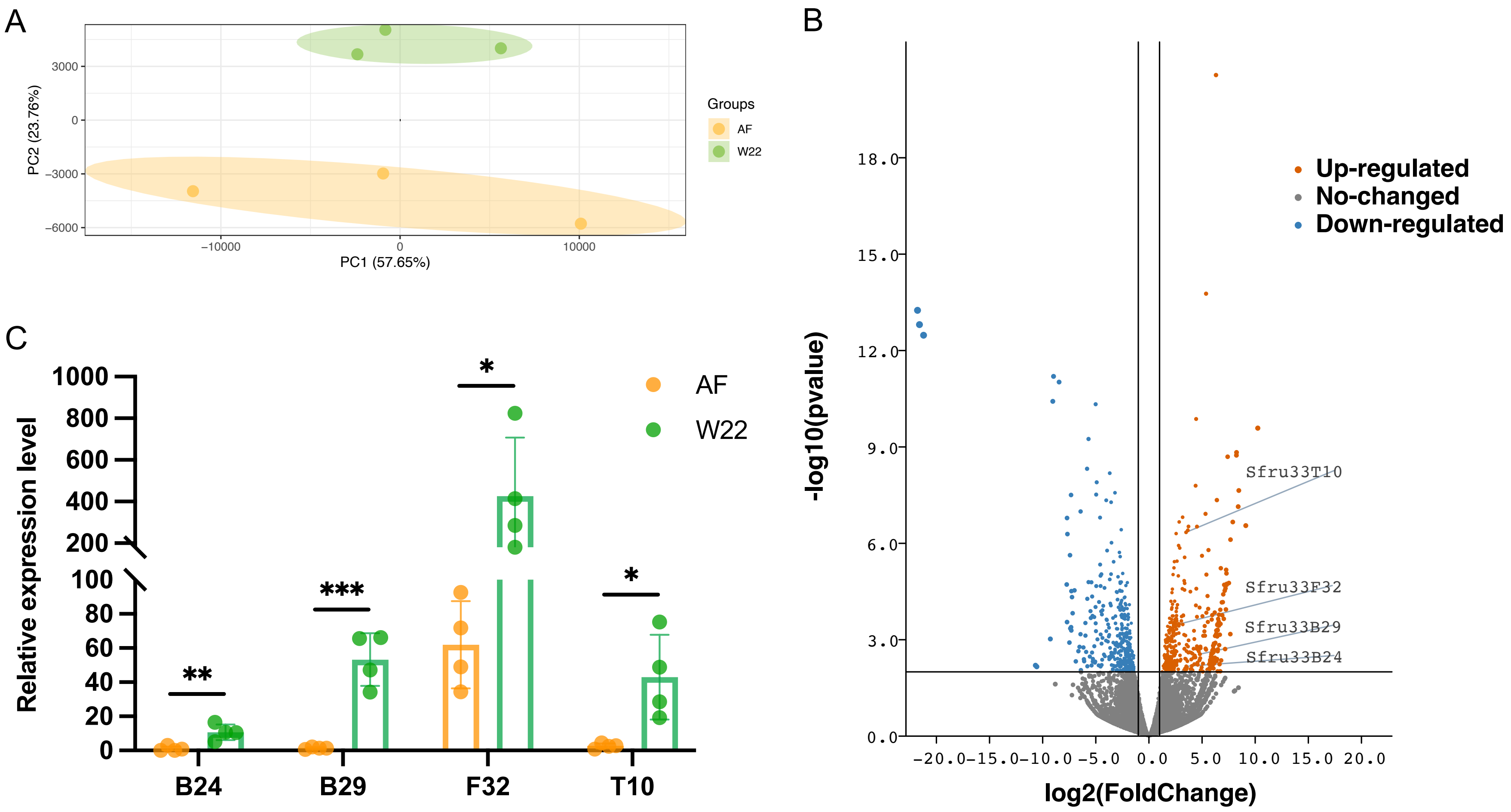


**Fig. S2. Transcriptomic analysis and differential gene expression of *S. frugiperda* feeding on wild-type maize (W22) or artificial food (AF).** (A) Principal component analysis (PCA) of RNA-seq data from the midguts of *S. frugiperda* fed on artificial food (AF) or wild-type maize (W22). (B) Volcano plot showing differential gene expression between the maize (W22) and AF treatments. The x-axis represents log2-fold change in gene expression, and the y-axis represents statistical significance (-log10 [p-value]). Genes with p < 0.05 and |fold change| > 1 were considered significantly differentially expressed. Red points indicate upregulated genes, blue points indicate downregulated genes, and gray points represent nonsignificant genes. The four candidate upregulated UGT genes are highlighted in the plot. (C) The relative expression of four candidate genes was confirmed by quantitative PCR (n = 4). Differences between the two groups (W22 and AF) were compared using Student's *t* test , and *, **, and *** represent $p < 0.05$, $p < 0.01$, and $p < 0.001$, respectively. Error bars represent standard error of the mean (SD).

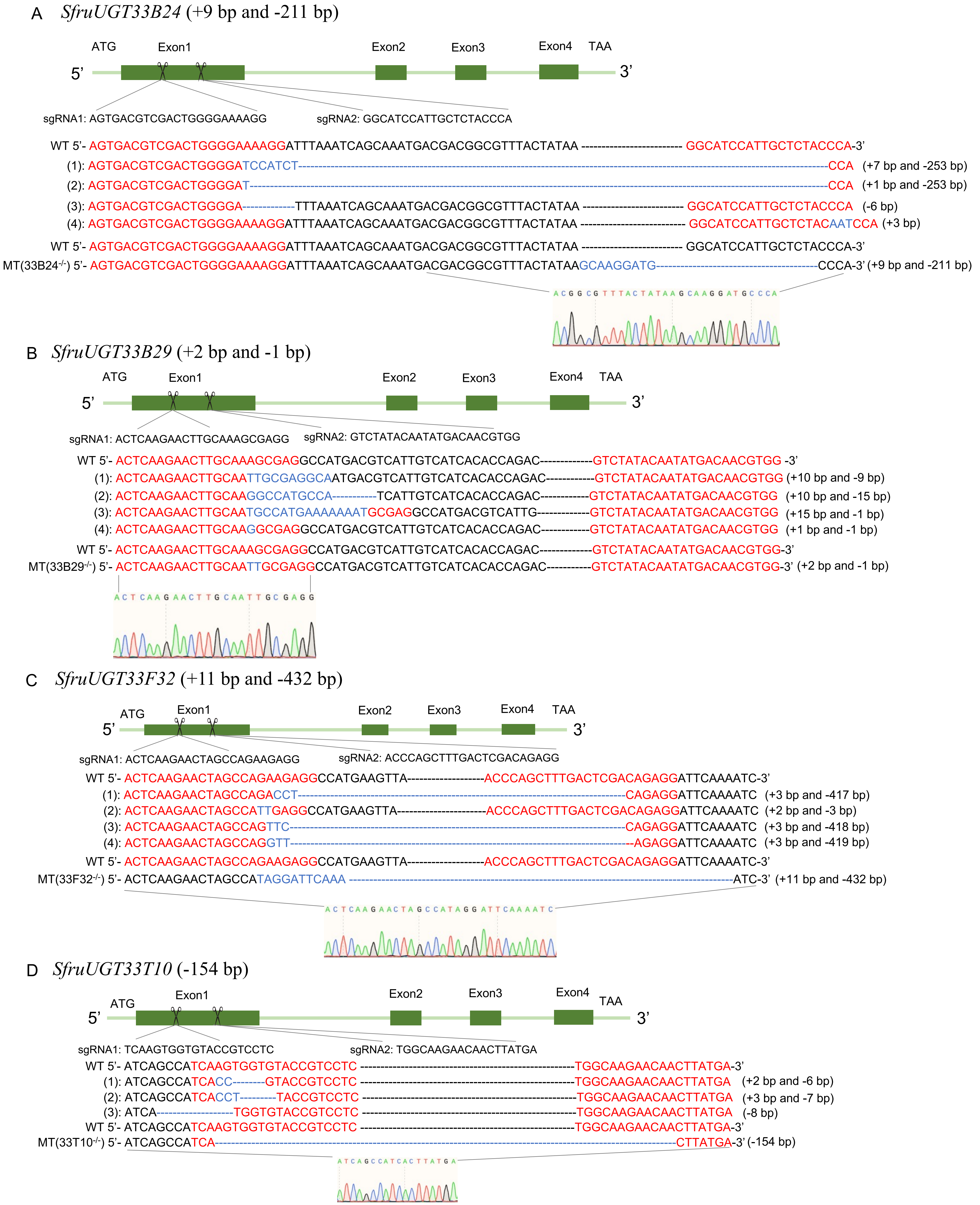


**Fig. S3. Schematic diagrams of CRISPR/Cas9-generated UGT mutants in *S. frugiperda*.** (A) *Sfru33B24*; (B) *Sfru33B29*; (C) *Sfru33F32*; (D) *Sfru33T10*. The gene sequence, target site, gene knockout status and sequencing results are shown in the figure. Each gene consists of four exons, with two knockout sites in exon 1. The mutations were generated using CRISPR/Cas9 with sgRNAs targeting exon 1, and confirmed by Sanger sequencing. Multiple mutation types were generated and confirmed by Sanger sequencing, including small insertions and deletions. Except for SfruUGT33B29, which carries small indels (+2 bp and -1 bp), the other mutants feature large fragment deletions. The final retained mutant lines are shown with homozygous mutations. In the sequences of mutants, red text indicates the sgRNA target sequences and blue text highlights edited regions. Each gene's stop codon appears within 20 bp after the knockout site. Sequencing chromatograms for the final mutants are shown.

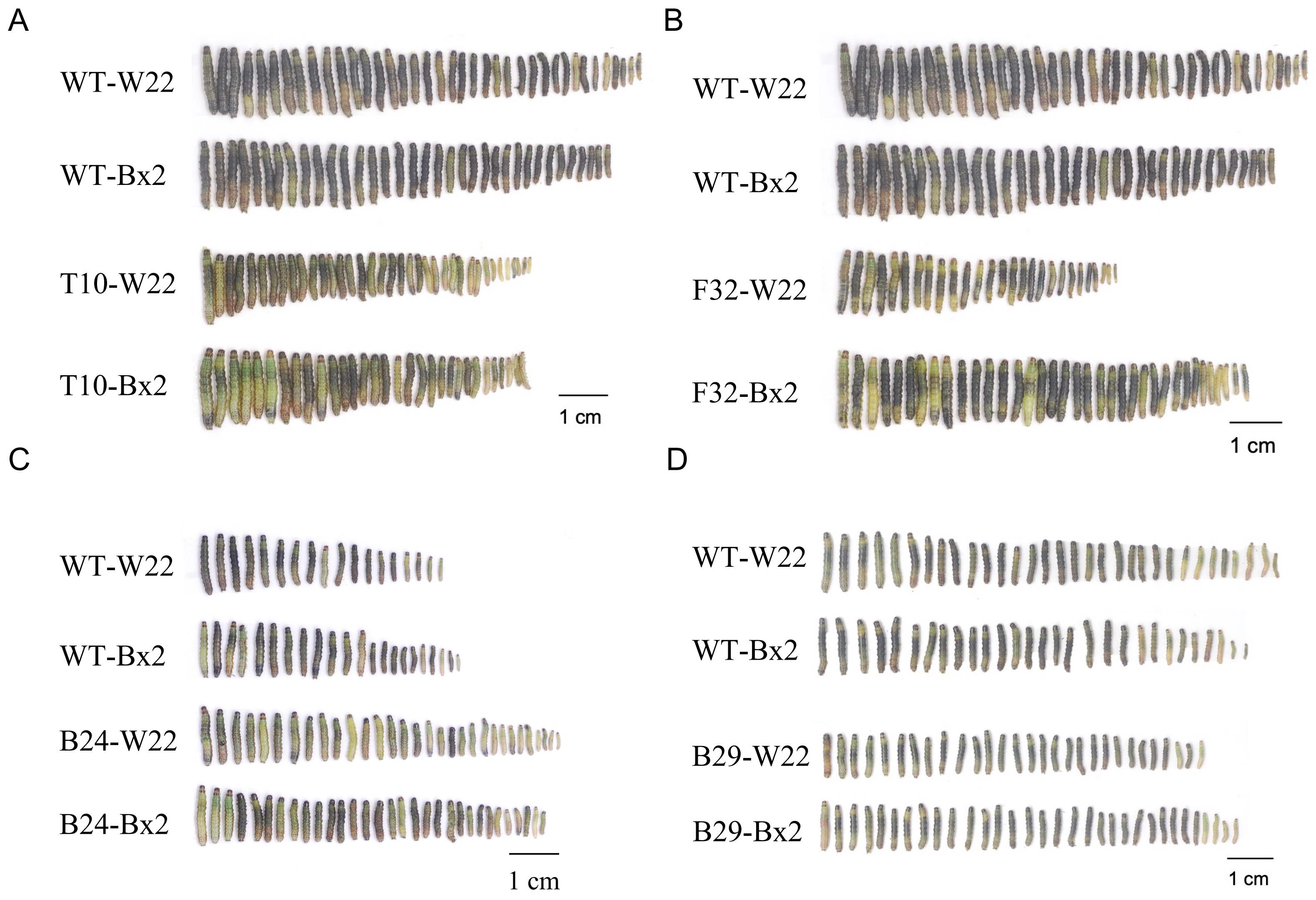


**Fig. S4. Photographs of different mutant larvae after feeding on maize.** (A) *Sfru33T10* mutant; (B) *Sfru33F32* mutant; (C) *Sfru33B24* mutant and (D) *Sfru33B29* mutant.

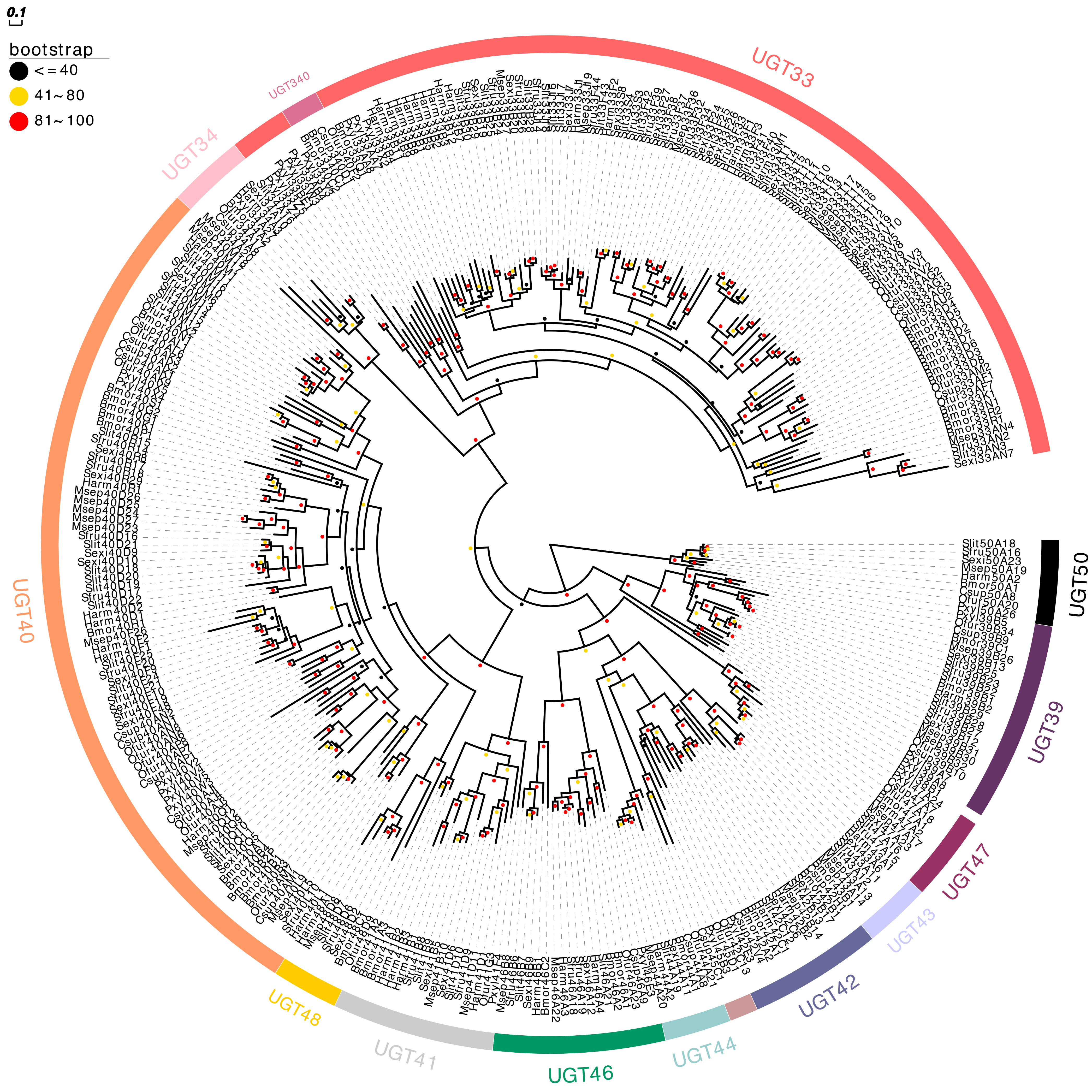


**Fig. S5. Phylogeny relationship of UGTs from nine Lepidoptera species.** UGT sequences from six species were annotated using de novo genome annotation. Sequences from *S. frugiperda, S. litura, S. exigua*, *M. separata, O. furnacalis*, and *P. xylostella* were included, while sequences from *H. armigera*, *B. mori* and *C. suppressalis* were obtained from published databases. The phylogenetic tree was constructed using maximum likelihood with 1000 bootstrap replicates and LG modol. Bootstrap values are represented by circles at branch nodes, with the size and color of circles indicating support levels. Leaf colors indicate UGT family groupings.

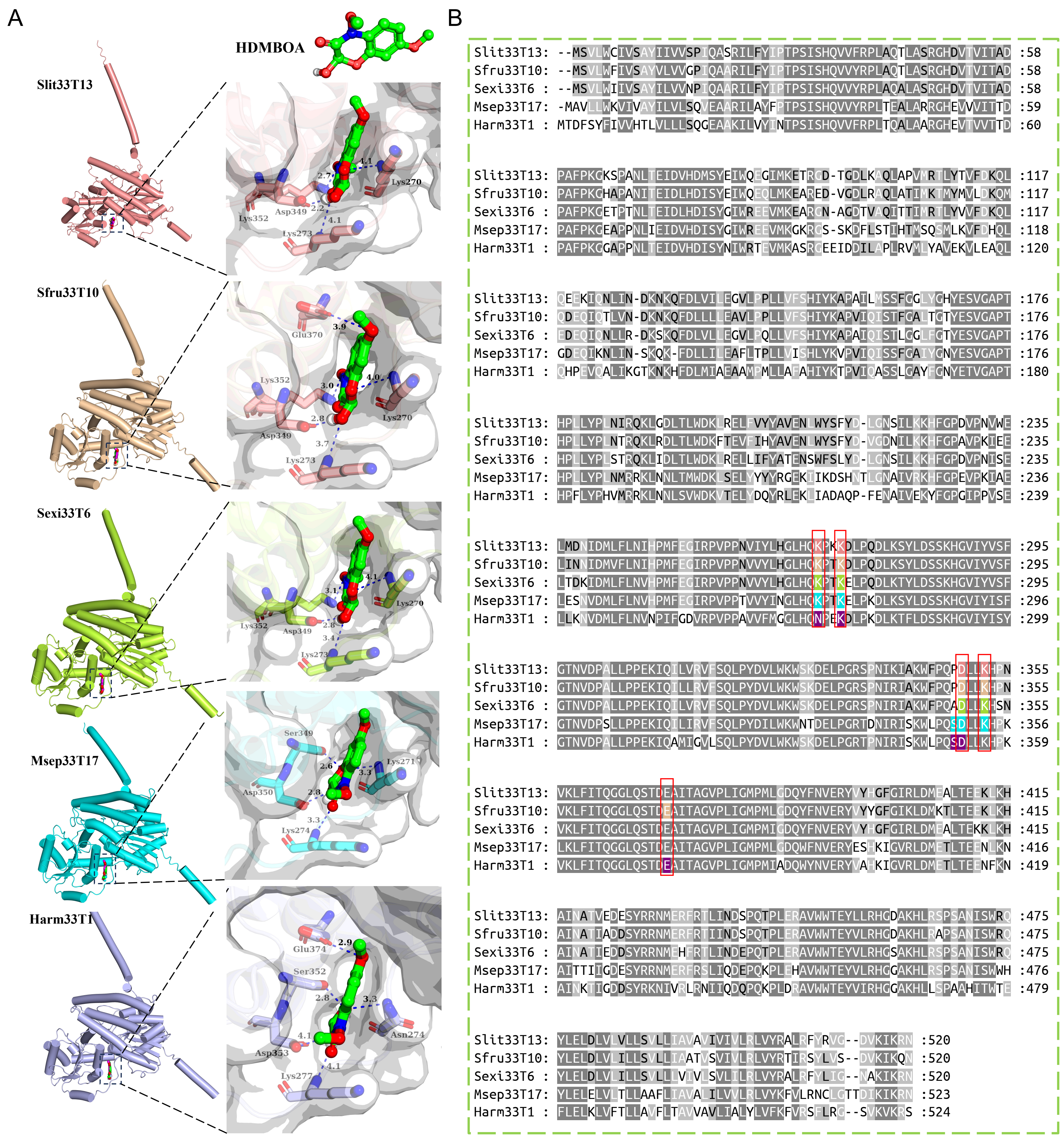


**Fig. S6. UGT33T proteins exhibit conserved binding sites for HDMBOA in five noctuid species.** (A) Molecular docking of HDMBOA and UGT33T proteins of five noctuid species. The binding pockets of the UGT33T proteins (*Sfru33T10*, *Slit33T13-SlitUGT33T13*, *Sexi33T6-SexiUGT33T6*, *Msep33T17-MsepUGT33T17*, and *Harm33T1-HarmUGT33T1*) are shown as gray surface, with interacting residues highlighted. Molecular docking was performed using AutoDock 4.2.6, and protein structures and interacting amino acid residues are displayed in different colors for each species: salmon for *SlitUGT33T13*, wheat for *SfruUGT33T10*, limon for *SexiUGT33T6*, cyan for *MsepUGT33T17*, and light blue for *HarmUGT33T1*. (B) Multiple sequence alignment of *SfruUGT33T10* and its homologs from five Noctuidae species. Key conserved amino acid residues are highlighted in red boxes, with different genes distinguished by various colors.

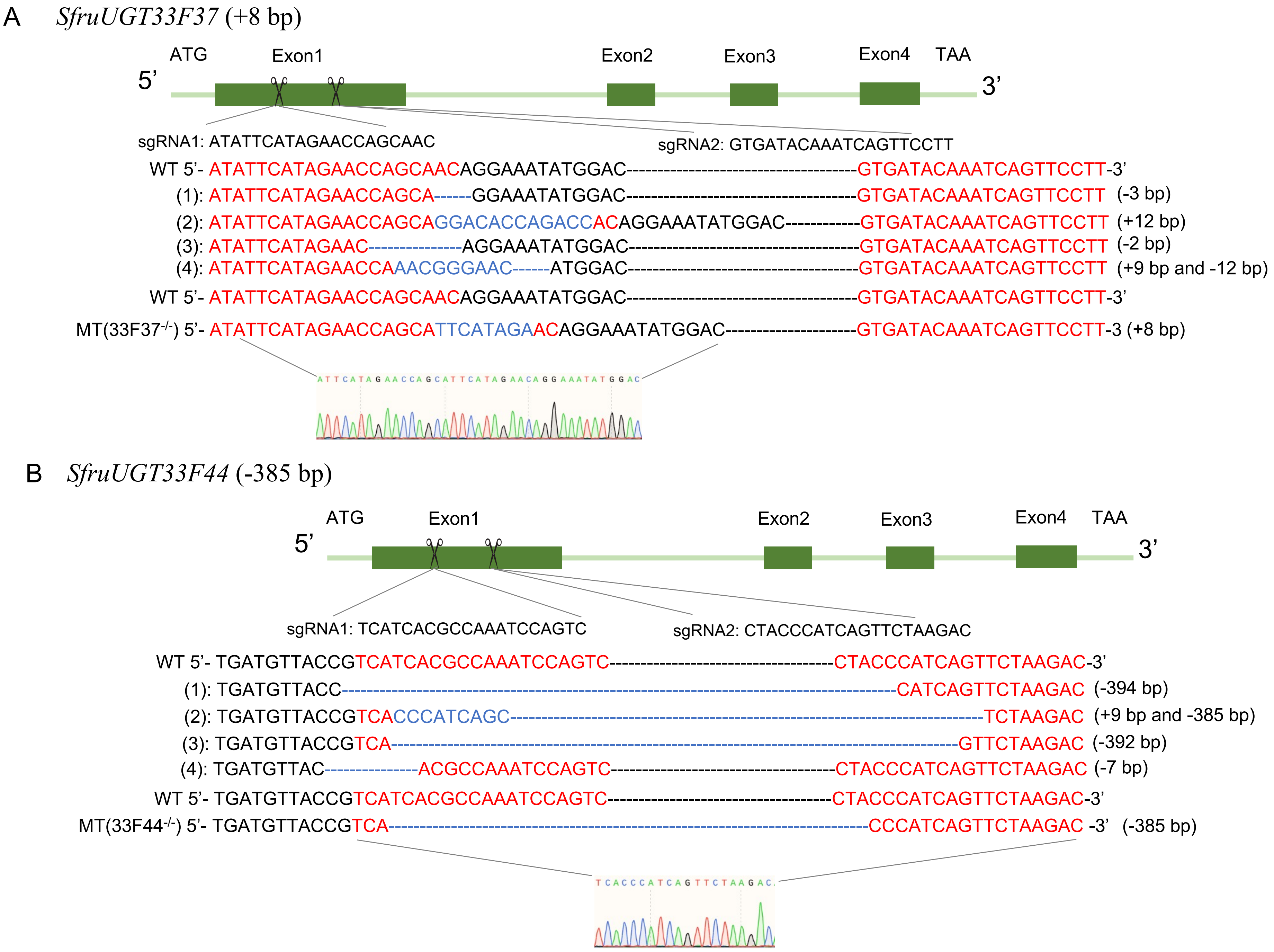


**Fig. S7. Schematic diagram of *Sfru33F37* and *Sfru33F44* mutants.** (A) *Sfru33F37*; (B) *Sfru33F44*. The gene sequence, target site, gene knockout status and sequencing results are shown in the figure. Each gene consists of four exons, with two knockout sites in exon 1. The mutations were generated using CRISPR/Cas9 with sgRNAs targeting exon 1, and confirmed by Sanger sequencing. Multiple mutation types were generated and confirmed by Sanger sequencing, including small insertions and deletions. The final retained mutant lines are shown with homozygous mutations. In the sequences of mutants, red text indicates the sgRNA target sequences and blue text highlights edited regions. Each gene's stop codon appears within 20 bp after the knockout site. Sequencing chromatograms for the final mutants are shown.
